# Mapping genetic modifiers of epimutation rates reveals a punctuated-equilibrium model of CG methylome evolution

**DOI:** 10.1101/2025.06.14.659605

**Authors:** Zhilin Zhang, Wilma Wanney, Yangyang Xu, Johan Zicola, Angela M. Hancock, Robert J. Schmitz, Frank Johannes

## Abstract

Spontaneous epimutations—stochastic changes in cytosine methylation—can persist across generations in plants and are thought to contribute to phenotypic variation. Although epimutations are increasingly studied for their potential long-term effects, it remains unclear why their accumulation varies across genotypes. Here, we tracked DNA methylation across ten generations in ∼400 mutation accumulation lineages derived from ∼70 *Arabidopsis* Ler x Cvi recombinant inbred lines. Treating epimutation rates as quantitative molecular traits, we mapped a major QTL to a Cvi-derived deletion near *VIM2* and *VIM4*, two genes involved in CG methylation (mCG) maintenance. We show that this deletion rapidly reduces genome-wide methylation to a lower steady-state and compromises mCG maintenance fidelity across generations, resulting in a ∼1.5-fold increase in epimutation rates. Genotypes with elevated rates exhibited accelerated epigenetic drift and phenotypic divergence. Our findings support a punctuated-equilibrium model of mCG evolution, in which sudden disruptions to methylation homeostasis can destabilize epigenetic inheritance over longer time-scales.

## Main

DNA cytosine methylation is a key epigenetic modification in plants, essential for transposable element (TE) silencing and gene expression ^1^. In both natural and domesticated plant populations, substantial intraspecific variation in methylation patterns has been documented ^2–6^, occasionally associated with stable, heritable phenotypic traits ^6–8^. In plants, cytosine methylation occurs in three sequence contexts—CG, CHG, and CHH (H = A, T, or C) ^9,10^ and is maintained by distinct pathways. CG methylation is preserved through DNA replication by METHYLTRANSFERASE 1 (MET1), in conjunction with VARIANT IN METHYLATION (VIM) proteins, which recognize hemimethylated CG sites and restore full methylation ^1,11–13^. In contrast, CHG and CHH methylation are maintained by CHROMOMETHYLASE 3 (CMT3), CMT2 and the *de novo* RNA-directed DNA methylation (RdDM) pathway ^1^.

Genome-wide association studies (GWAS) have shown that natural variation in non-CG methylation is frequently associated with methylation quantitative trait loci (meQTLs) ^2,4,14–16^. These loci typically map to core RdDM or chromatin remodeling components, and associated alleles often produce pleiotropic effects on TE methylation throughout the genome ^16^. Such findings point to a relatively direct genetic basis for non-CG methylation variation, rooted in perturbations of known regulatory networks. In contrast, CG methylation (mCG) presents a more complex pattern. Although mCG levels—defined as the proportion of methylated CG sites in a genome—vary widely among *A. thaliana* accessions ^4^, their genetic basis remains elusive. A likely reason is the high frequency of stochastic methylation gains and losses (i.e., spontaneous epimutations) that also accumulate at CG sites over evolutionary time scales ^17,18^. These stochastic events result from maintenance errors, and are often stably inherited across generations ^10,19^. This property allows mCG to act both as a molecular trait and as a heritable state, capable of segregating independently of genetic variation^20^.

Spontaneous epimutations occur at a rate of ∼10^-4^ per site per generation—about 100,000 times higher than the DNA mutation rate ^9,21–25^. Theory predicts that the continuous interplay between spontaneous methylation gain and loss drives mCG levels toward a dynamic equilibrium, with a steady-state determined by their relative rates ^9,26,27^. Observed methylation levels across diverse *A. thaliana* accessions are broadly consistent with this prediction ^21^. However, some accessions exhibit unusually high or low mCG levels relative to the population average ^4^. These outliers may reflect alternative steady-states driven by shifts in the underlying epimutation dynamics—potentially caused by genetic variants that alter the fidelity of methylation maintenance across generations. Testing this hypothesis requires a system capable of jointly mapping both steady-state methylation and epimutation dynamics—an approach not possible with cross-sectional GWAS. Natural mCG outliers provide a valuable opportunity to explore this question and to uncover genetic variants that influence epigenetic stability.

To address this gap, we focused on Cvi, an *A. thaliana* accession with globally reduced mCG levels, and generated a large population of mutation accumulation mapping lines (MAML). This population was derived from ∼70 Ler × Cvi recombinant inbred lines (RILs) founders, propagated over multiple generations to produce ∼400 independently evolving mutation accumulation (MA) lineages. Using this system, we identified a major QTL on chromosome 1 with strong effects on both mCG levels and epimutation rates. The causal variant is a Cvi-specific structural deletion that disrupts *VIM2/4* expression and its co-regulation with other methylation and demethylation pathway genes. We show that this QTL induces a rapid reduction in mCG levels, followed by elevated methylation gain and loss rates, leading to accelerated epigenetic drift and functional divergence within genotypes over time. These findings link genetically driven shifts in methylation to long-term instability of the methylome, revealing how deterministic and stochastic processes interact in epigenome evolution.

## Results

### Genome-wide CG methylation levels and epimutation rates are genotype dependent

To quantify natural variation in mCG, we re-analyzed published DNA methylomes from 850 *A. thaliana* accessions ^4^. Genome-wide mCG levels were ∼2.5-fold more variable among accessions than non-CG methylation (Fig. 1a and Supplementary Table 1), emphasizing the relevance of this context. To assess genotype-specific differences in mCG stability across generations, we previously established mutation accumulation (MA) pedigrees for four selected accessions (Col-0, Kn-0, Mt-0, Tsu-0) that span the natural CG methylation range, and propagated them for up to 16 selfings^21^ (Fig. 1a,b). Epimutation analysis revealed up to 1.4-fold differences in spontaneous mCG gain (α) and loss (β) rates, indicating that some genotypes are more error prone at maintaining CG methylation states transgenerationally (Fig. 1c,d). Predicted mCG levels based on these rates closely matched empirical observations in each accession (Fig. 1e), implying that all genotypes are at methylation equilibrium ^9,26^. These findings demonstrate that steady-state mCG levels and epimutation rates are determined, in part, by genetic background.

**Fig. 1.**
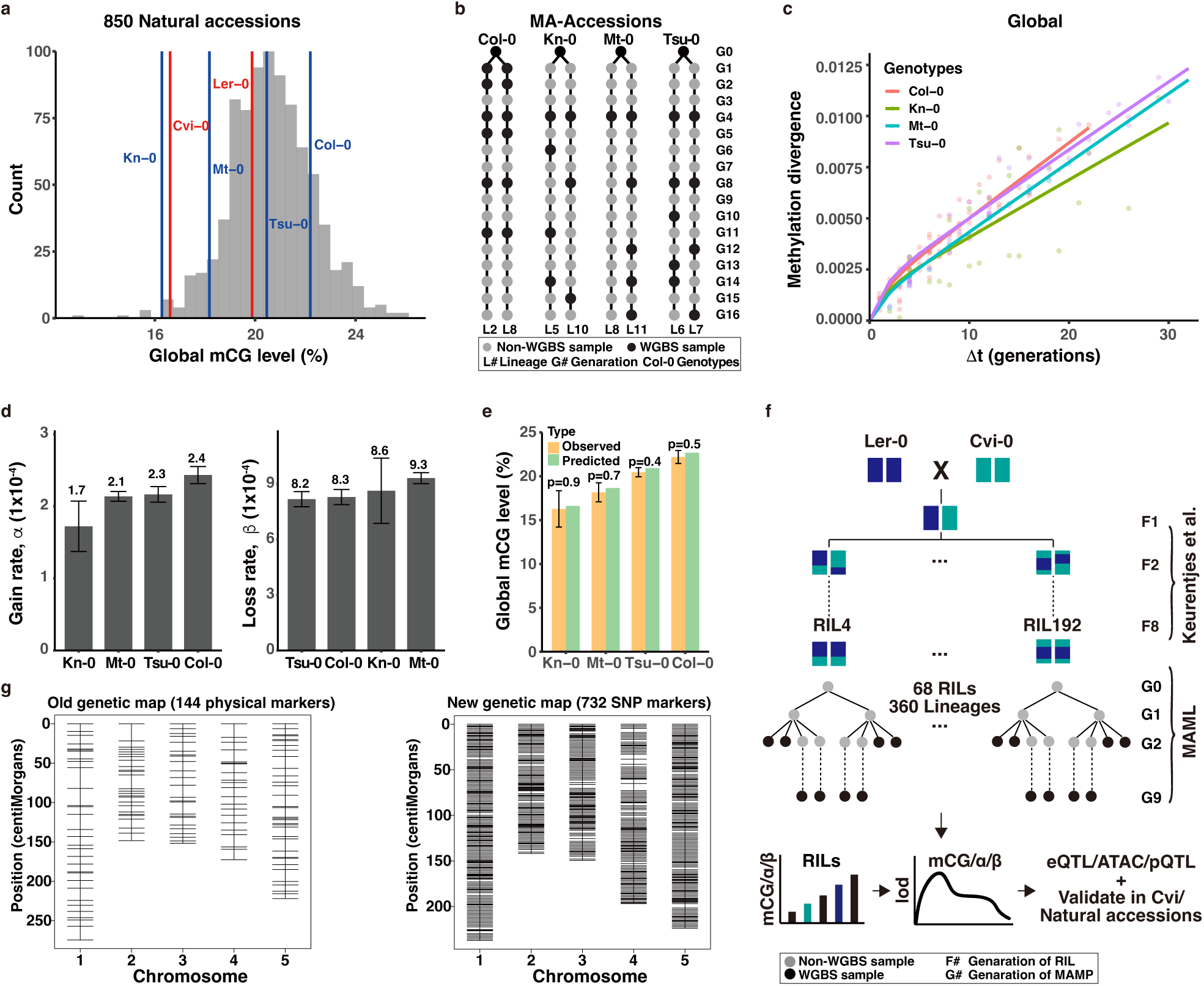
Genotype-dependent methylation variation and the establishment of a mutation accumulation mapping lines (MAML) population. **a**, Distribution of genome-wide CG methylation (mCG) levels across 850 natural *Arabidopsis thaliana* accessions. Blue lines represent accessions selected for mutation accumulation (MA) experiments (Col-0, Kn-0, Mt-0, Tsu-0) and red lines indicate the Ler-0 and Cvi-0 lines used in the MAML population. **b**, Schematic of MA pedigrees established for four selected accessions spanning the natural CG methylation range, propagated for up to 16 generations by single-seed descent. Black circles indicate samples sequenced by whole-genome bisulfite sequencing (WGBS). **c**, Global methylation divergence as a function of generational time (Δt, generations) in four MA lines, illustrating genotype-specific differences in mCG stability. Curves represent model fits for each genotype. **d**, Comparison of genome-wide spontaneous mCG gain rates (α, left panel) and loss rates (β, right panel) among the four genotypes. Error bars denote standard error of the mean (SEM). **e**, Observed versus predicted genome-wide CG methylation levels based on epimutation rates, indicating that genotypes are at methylation equilibrium. Error bars represent SEM; p-values from one-sample t-tests comparing observed and predicted values are indicated above bars. **f**, Overview of the experimental framework used to dissect the genetic basis of methylation dynamics. The MAML were constructed from Ler × Cvi recombinant inbred lines (RILs). Each RIL was used to initiate eight independent MA lineages, propagated for ten generations (single-seed descent), resulting in 68 MAML pedigrees. Additional MA lines were also derived from the Ler-0 and Cvi-0 parental accessions. WGBS datasets were generated from early (G2) and late (G9) generations to quantify mCG changes over time, enabling subsequent QTL analyses. These data were integrated with previously published expression QTL (eQTL) and phenotypic QTL (pQTL) resources derived from Ler × Cvi RILs, along with newly generated chromatin accessibility profiles from assay for transposase-accessible chromatin using sequencing (ATAC-seq), to support multi-layered trait dissection and were further validated in Cvi and natural accessions. **g**, Comparison of the original genetic map (144 physical markers, left panel) used in previous studies versus the newly constructed genetic map (732 SNP markers, right panel) generated from SNPs called from WGBS data in this study, substantially enhancing mapping resolution.

### Construction of Mutation Accumulation Mapping Lines (MAML) population

To identify genetic factors contributing to variation in mCG maintenance dynamics, we generated the MAML population using Landsberg erecta (Ler-0) × Cape Verde Islands (Cvi-0) RILs as founders ^28^. Each RIL was used to initiate eight independent MA lineages, propagated by single-seed descent for ten generations (Fig. 1f). The Ler x Cvi RILs were chosen because their parental accessions have highly contrasting mCG levels, with Cvi-0 scoring in the first percentile of the natural distribution and Ler-0 ranking substantially higher (Fig. 1a). We expected that this wide cross maximizes our ability to detect genetic variations underlying mCG differences. Moreover, the Ler × Cvi RILs have been extensively characterized in prior QTL studies of both classical and molecular phenotypes, including transcript, metabolite, and protein abundance ^29,30^, thus providing a rich framework for integrative analysis. To capture methylation dynamics across generations, we performed whole-genome bisulfite sequencing (WGBS) on early- and late-generation individuals from 68 MAML pedigrees as well as from MA lines derived from the Ler-0 and Cvi-0 parental accessions, yielding 371 WGBS datasets in total (Fig. 1f and Supplementary Table 2). Using these data, we saturated the existing genetic map with additional single-nucleotide polymorphisms (SNPs) called from the WGBS reads (Methods). This resulted in a genetic map comprising 732 SNP markers, with an average spacing of 1.3 cM and a maximum interval of 8.0 cM—substantially improving upon the previously published map (Fig. 1g, Supplementary Fig. 1, and Supplementary Tables 3 and 4).

### Mapping genetic mediators of methylation levels

Genome-wide DNA methylation profiling of MAML revealed a 1.4-fold variation in global mCG levels across genotypes, ranging from 14.7% to 21.3%. The parental accessions Ler-0 and Cvi-0 scored toward the extremes of this distribution (Fig. 2a), consistent with limited transgressive segregation ^31^. Comparisons of mCG levels between G2 individuals and their G9 descendants for each genotype revealed no significant differences (Fig. 2b), indicating that each lineage had reached mCG steady-state already at the onset of MAML construction. This stability contrasts with the progressive methylation loss typically observed in many DNA methylation mutants ^32–34^. Based on this, we pooled genome-wide mCG data from all samples within each pedigree. Treating mCG variation across genotypes as a quantitative molecular trait, we performed linkage-based methylation QTL (meQTL) mapping. This analysis identified a single major-effect meQTL on chromosome 1 that accounted for 44% of the variance in global mCG (LOD = 8.62; Confidence Interval (CI): 23.7–25.8 Mb) (Fig. 2c and Supplementary Table 5). To explore whether additional annotation-specific effects were masked in the global analysis, we performed stratified mapping across distinct genomic features. The chromosome 1 meQTL exhibited its strongest effect on mCG levels within gene body methylated (gbM) genes, explaining approximately 59% of the variance (LOD = 13.11; CI: 24.6–25.7 Mb), and also revealed two minor-effect loci on chromosomes 3 and 5. In addition, a distinct, strong-effect meQTL was detected at the distal end of chromosome 2, specifically influencing mCG levels in TEs, TE-like methylated (teM) genes, and intergenic regions, accounting for 23%, 30%, and 26% of the variance in these features, respectively (Fig. 2c and Supplementary Table 5).

**Fig. 2.**
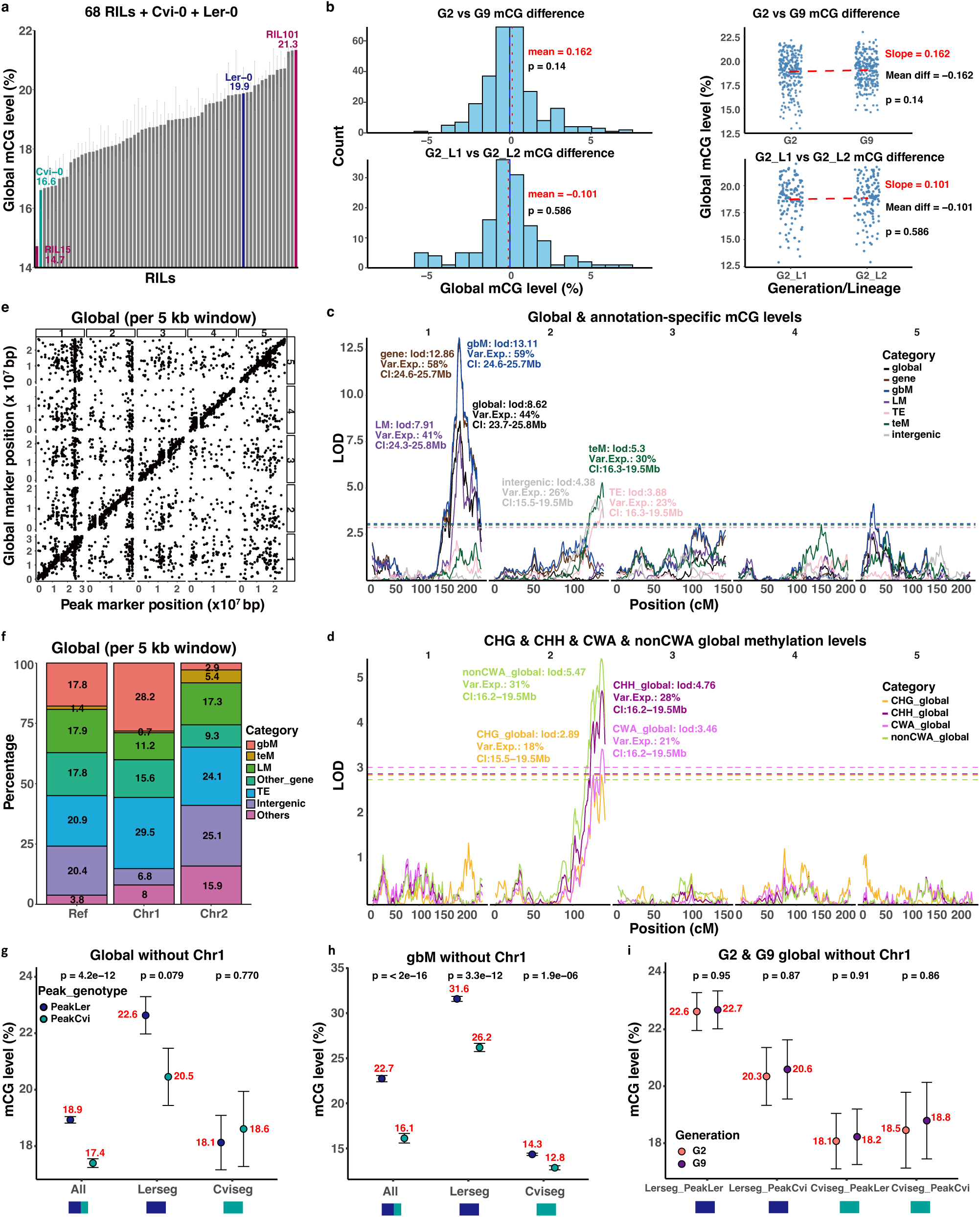
Mapping genetic mediators of methylation levels. **a**, Genome-wide mCG levels across 68 RILs derived from Ler × Cvi and their parental lines. Lines are ranked by the mean mCG level averaged across all samples for each RIL. Dark magenta labels indicate RILs with extreme mCG levels. The parental lines, Ler-0 and Cvi-0, are highlighted in blue and teal, respectively. **b**, Stability of mCG levels within RILs across generations and lineages. Top: histograms (left) and paired plots (right) show negligible differences between early (G2) and late (G9) generations within each RIL, indicating steady-state methylation levels. Bottom: comparisons between two independent lineages of the same RIL show similarly high concordance in mCG levels. Paired t-tests were used for both comparisons; p-values are indicated. **c**, Methylation QTL (meQTL) mapping results using global and annotation-specific mCG levels as quantitative traits. A major-effect QTL on chromosome 1 significantly affects global, gene, and gbM methylation levels, while an additional locus on chromosome 2 specifically influences methylation in TE, teM, and intergenic regions. Peak LOD scores (“LOD”), Explained Variance (“Var.Exp.”), and Confidence Intervals (“CI”) are indicated. Horizontal dashed lines represent significance thresholds determined by permutation testing (α = 0.05, 1000 permutations). Position is given in centiMorgans (cM). **d**, QTL mapping for global non-CG methylation identified a major-effect locus for CHG, CHH, CWA, and non-CWA methylation on chromosome 2, co-localizing with the mCG meQTL associated with TE, teM, and intergenic regions shown in (c). **e**, Genome-wide *cis–trans* analysis of mCG in 5 kb windows reveals two major *trans*-acting QTL bands co-localizing with chromosome 1 and chromosome 2 meQTL. **f**, Genomic feature enrichment within *trans*-bands highlights distinct target preferences of chromosome 1 (enriched for gbM genes) and chromosome 2 (enriched for TEs, teM, and intergenic regions) meQTL. **g-h**, Haplotype-specific and directional effects of the chromosome 1 meQTL on global (**g**) and gbM-specific (**h**) CG methylation. The Cvi allele at the peak of the chromosome 1 meQTL (PeakCvi) significantly reduces methylation levels in Ler-derived genomic segments (Lerseg), with little effect on Cvi-derived segments (Cviseg). Error bars represent SEM; p-values are from two-sided t-tests. **i**, Global mCG levels in G2 and G9 generations across genotypes stratified by chromosome 1 meQTL haplotype, showing that the genetic effect is established early and remains stable across generations.

Given the frequent co-regulation of mCG with non-CG methylation (mCHG and mCHH) in these regions through shared epigenetic pathways ^1,35^, we hypothesized that the chromosome 2 meQTL might also affect non-CG methylation. Supporting this, mapping of global mCHG and mCHH levels identified a major-effect locus that co-localized with the chromosome 2 mCG meQTL. The strongest association was observed for CHH methylation, particularly at non-CWA (CWA refers to CAA or CTA) contexts (Fig. 2d and Supplementary Table 5), hinting at an involvement of the RdDM pathway. Interestingly, *FBX5* emerged as a significant *cis*-eQTL (expression QTL) target located within the QTL confidence interval in our analysis (Supplementary Table 6), consistent with its recent identification by Zicola et al. as a novel modifier of DNA methylation in natural Cvi accessions (see co-submission).

The target-specificity of the two major meQTL was further supported by repeating the analysis using chromatin state (CS) annotations of the *A. thaliana* genome ^36,37^. The chromosome 2 locus was specifically associated with four CSs enriched in intergenic and TE regions, while the chromosome 1 locus was predominantly associated with mCG levels in CS characteristic of gbM subregions ^36^ (Extended Data Fig. 1 and Supplementary Table 7).

The strong effects of the chromosome 1 and chromosome 2 meQTL on both global and feature-specific mCG levels suggested that they act primarily in *trans* to modulate methylation at distal targets. To test this directly, we conducted a genome-wide *cis–trans* analysis by segmenting the genome into 5-kb windows and treated regional mCG levels as quantitative traits. This revealed two prominent *trans*-bands co-localizing with chromosomes 1 and 2 meQTL (Fig. 2e). As expected, the chromosome 1 *trans*-band was enriched for gbM, whereas the chromosome 2 *trans*-band was enriched for TEs, teM genes, and intergenic regions (Fig. 2f), consistent with prior annotation-stratified results (Fig. 2c and Extended Data Fig. 1). We examined the directionality and haplotype specificity of these *trans*-acting effects in detail. We found that the Cvi allele at the chromosome 1 locus reduced mCG levels at distal targets to levels approximating those of the Cvi-0 parent—but only within Ler-derived genomic segments (Fig. 2g,h). This suggests that the Cvi allele induces hypomethylation primarily at pre-methylated Ler-inherited loci. To confirm the transgenerational stability of genetic effects on mCG levels, we independently performed meQTL mapping in the G2 and G9 generations. Both analyses revealed the same genetic architecture underlying the hypomethylation phenotypes, indicating that these effects are established early and persist across generations (Fig. 2i and Extended Data Fig. 2a).

Together, these findings demonstrate that the meQTL elicits rapid and stable transitions to new methylation steady-states in MAML genomes, primarily by acting in *trans* to reduce methylation on Ler-derived genomic segments.

### A major QTL modulates the spontaneous CG epimutation rates

We asked whether genetically driven perturbations in steady-state CG methylation can have secondary effects on the fidelity of mCG maintenance across generations. Prior work in *CMT3* mutants demonstrated that even modest changes in global methylation can compromise epigenetic stability, increasing susceptibility to the accumulation of spontaneous epimutations over generations—even when steady-state levels appear robust ^36^ (Methods). To test this in the MAML population, we estimated genotype-specific epimutation rates across all pedigrees using established tools ^9,22^. Rates varied widely among RILs, with the most hyper-epimutable line reaching 14.5 x 10^-4^ and the most stable line 3.3 x 10^-4^ per site per haploid genome per generation—a 4.4-fold difference in epigenomic fidelity (Fig. 3a). These differences were mirrored in the rate of mCG divergence over generations, with hyper-epimutable lines showing markedly faster epigenetic drift (Fig. 3b). These results align with findings from our MA lines in natural accessions (Fig. 1c,d), and confirm that the fidelity of mCG maintenance is inherently genotype-dependent.

**Fig. 3.**
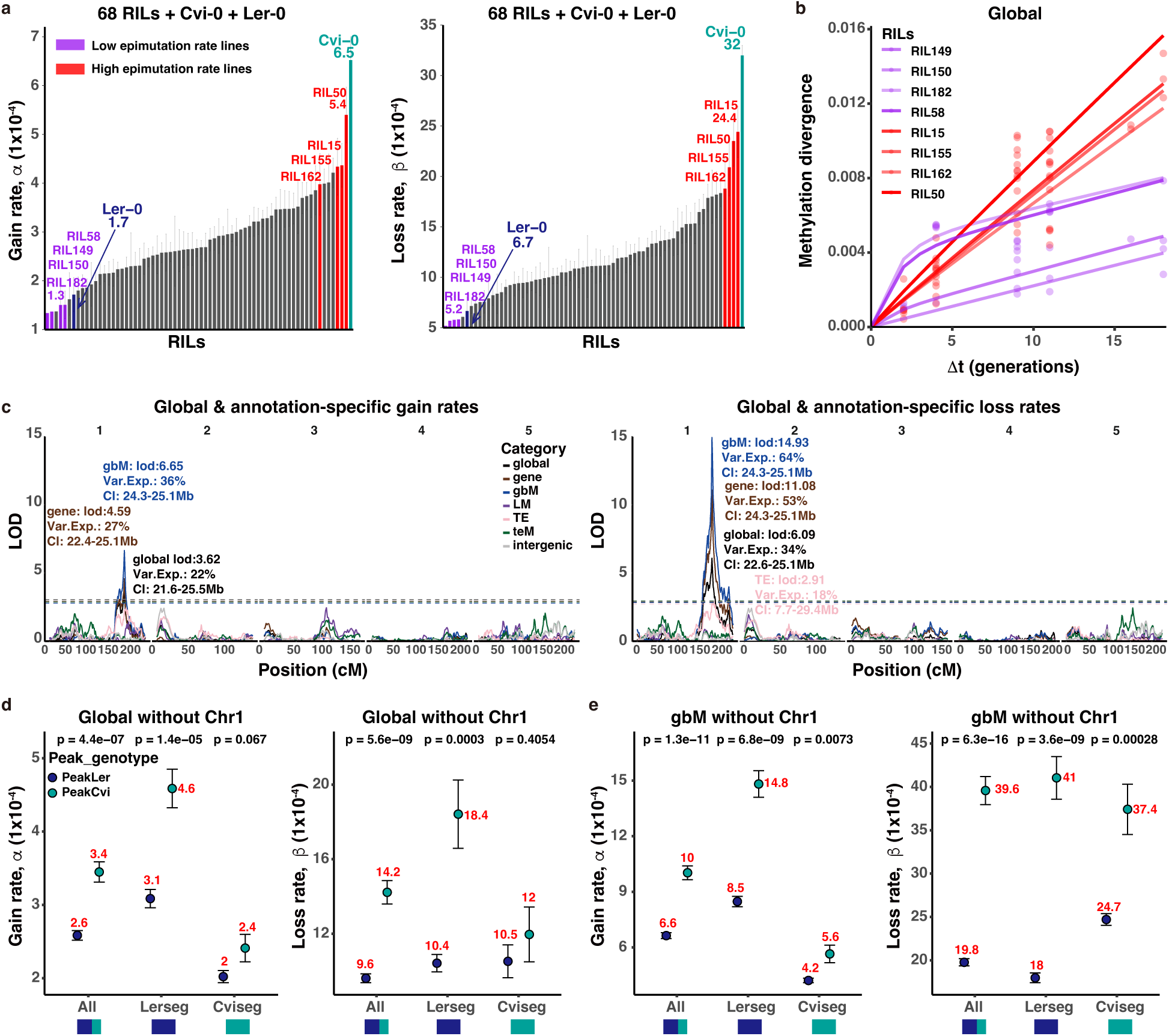
Mapping genetic modifiers of CG epimutation rates. **a**, Distribution of genome-wide estimates of spontaneous CG epimutation gain (left) and loss (right) rates across 68 RILs and their parental lines (Ler-0, Cvi-0). Lines are ranked by rate; RILs with extreme epimutation rates and parental lines are color-highlighted. **b**, Global methylation divergence over multiple generations for selected RILs with extreme epimutation rates, illustrating faster divergence in hyper-epimutable lines (red) compared to stable lines (purple). **c**, QTL mapping results for global and annotation-specific spontaneous CG methylation gain (left) and loss (right) rates. A major-effect locus on chromosome 1 significantly modulates both gain and loss rates, with a notably stronger effect on loss rates, particularly within gbM genes. Statistical metrics are labeled as in Fig. 2. This locus co-localizes with the previously identified meQTL for steady-state mCG levels. Other loci on chromosomes 2, 3, and 5 were not associated with epimutation rate variation. **d-e**, Haplotype-specific effects of the chromosome 1 QTL on global (**d**) and gbM-specific (**e**) epimutation rates. The Cvi allele (PeakCvi) increases both gain and loss rates predominantly in Ler-derived genomic regions (Lerseg), with minimal effects in Cvi-derived regions (Cviseg).

To dissect the genetic basis of this variability, we treated epimutation rates as molecular traits and performed QTL mapping. This analysis identified a single major-effect locus on chromosome 1 that influenced both gain and loss rates, with a notably stronger effect on loss rates. The QTL explained 22% of the variance in global CG gain rates and 34% in loss rates, with particularly pronounced effects in gbM genes, where it accounted for 36% and 64% of the variance, respectively (Fig. 3c, Extended Data Fig. 2b, and Supplementary Tables 8 and 9). This locus precisely co-localized with the previously mapped QTL for steady-state mCG levels. In contrast, the minor-effect QTL on chromosomes 3 and 5, as well as the major chromosome 2 locus associated with non-CG methylation (CHG/CHH) and TE/intergenic CG methylation, were not associated with variation in epimutation rates (Figs. 2c-f and 3c, Extended Data Figs. 1 and 2, and Supplementary Tables 5 and 7-9). This observation suggests that while these latter loci shape mCG levels, they do not influence transgenerational methylation dynamics. We previously argued that CG sites within or proximal to regions targeted by RdDM are less prone to accumulate epimutations ^9,36^, likely due to periodic reinforcement during gametogenesis and early embryogenesis. Thus, QTL that co-segregate with mCG levels may not necessarily influence long-term maintenance fidelity.

To explore the haplotype specificity of these effects, we examined epimutation patterns in Ler- and Cvi-derived chromosomal segments in more detail. We found that the elevated loss rates associated with the Cvi allele at the chromosome 1 QTL were largely restricted to Ler-derived genomic regions (Fig. 3d,e), consistent with earlier findings that the Cvi allele drives hypomethylation of these segments (Fig. 2g,h). Interestingly, Ler segments also displayed a higher propensity for methylation gain in the presence of the Cvi allele (Fig. 3d,e), pointing to a more general destabilization of mCG maintenance in these segments across generations.

Taken together, these observations highlight a model in which genetically driven methylation changes give rise to new epigenetic steady-states that are intrinsically less stable and more susceptible to error-prone maintenance.

### The chromosome 1 QTL perturbs CG methylation pathways via altered *VIM* regulation

To begin to understand the mechanistic basis of the chromosome 1 QTL, we searched for candidate genes using expression QTL (eQTL) mapping with previously generated transcriptome data from the Ler × Cvi RIL population ^29,30^. This analysis identified *VARIANT IN METHYLATION 2/4* (*VIM2* and *VIM4* combined expression) as a significant *cis*-eQTL target located within the QTL confidence interval (Fig. 4a and Supplementary Table 10). The *VIM* gene family encodes proteins that cooperate with MET1 to maintain CG methylation during DNA replication ^1,11–13^. *VIM4* lies ∼2.6 kb from *VIM2* (Fig. 4a), and as a paralog shares high sequence and structural similarity, suggesting potential co-regulation. A recent GWAS of mCG variation in natural Cvi accessions on the Cape Verde Islands also identified a strong association at this locus (Zicola et al., co - submission). Read coverage analysis revealed a ∼2.7 kb Cvi-specific deletion between *VIM2* and *VIM4*, which segregates in both natural Cvi accessions population and the MAML population (Fig. 4a). The deletion spans three small TEs: AT1TE80750 (∼100 bp, ATREP15), AT1TE80755 (∼1 kb, ATREP3), and AT1TE80760 (∼20 bp, ATREP15). Among these, AT1TE80755 is strongly methylated in the reference accessions Col-0 and Ler-0 (Fig. 4a), suggesting a potential role in transcriptional repression. Consistent with this idea, RILs carrying the Cvi haplotype, which lacks these TEs, showed significantly elevated *VIM2/4* expression compared to lines with the intact Ler haplotype (Fig. 4a,b). Hence, methylation at AT1TE80755 appears to function as a local repressive element that constrains *VIM* expression in accessions such as Ler-0.

**Fig. 4.**
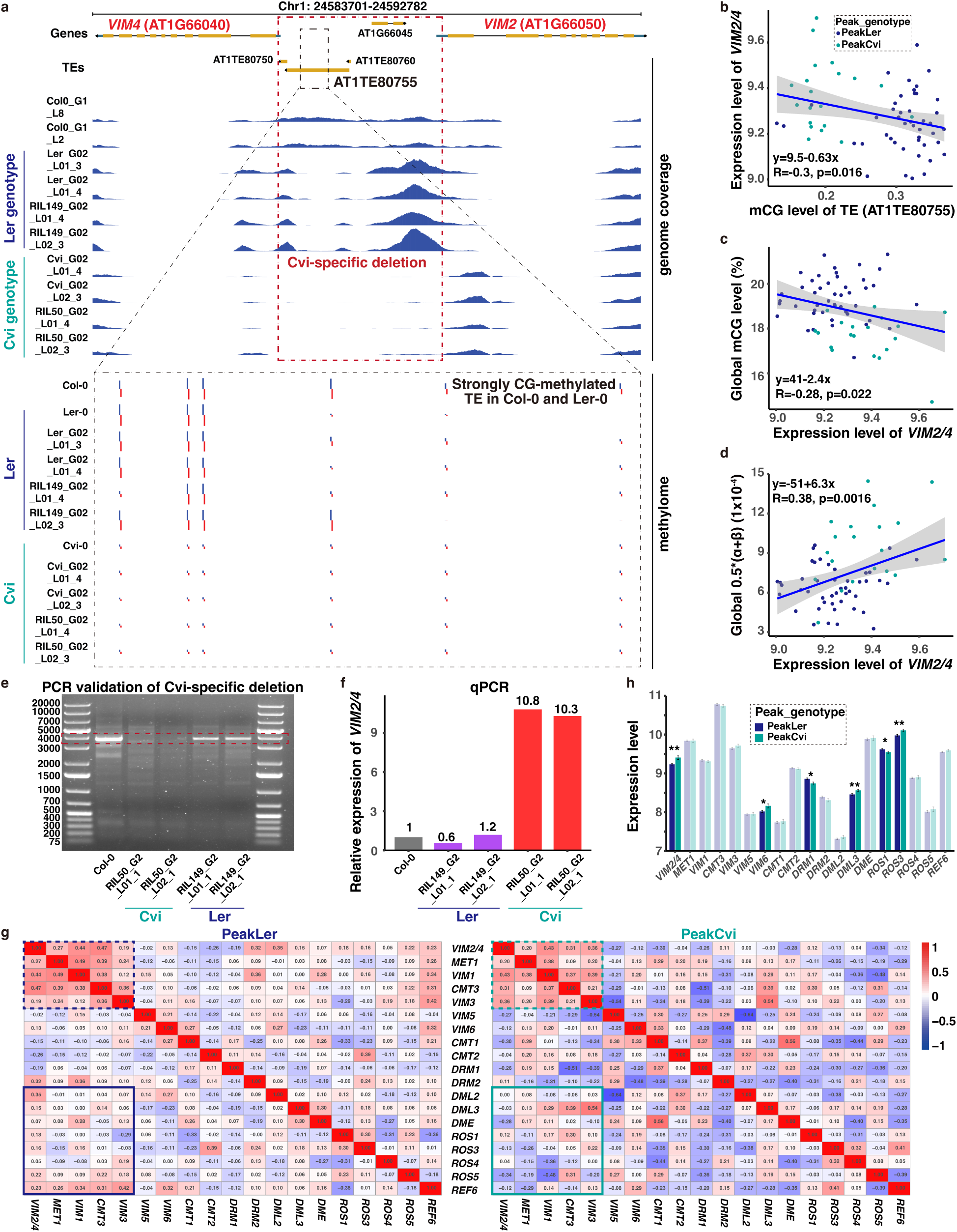
A Cvi-specific deletion between *VIM2* and *VIM4* perturbs CG methylation maintenance through enhanced *VIM* expression. **a**, Genomic view of the *VIM2–VIM4* region showing genome coverage (top) and DNA methylation (bottom) across two RILs with contrasting epimutation rates—RIL50 (hyper-epimutable; Cvi genotype at the Chr1 methylation QTL) and RIL149 (epigenetically stable; Ler genotype at the Chr1 methylation QTL), along with the Col-0, Ler-0, and Cvi-0 accessions lines, and the natural accessions (“Col-0”, “Ler-0”, and “Cvi-0”). Two biological replicates are shown for each RIL for DNA methylation and genome coverage to illustrate reproducibility. A ∼2.7 kb Cvi-specific deletion (dashed red box) between *VIM2* and *VIM4* removes three transposable elements (AT1TE80750, AT1TE80755, and AT1TE80760), including a strongly methylated ATREP3 element (AT1TE80755; dashed black box) in Ler-0 and Col-0. **b–d**, Relationships between combined *VIM2* and *VIM4* (i.e., *VIM2/4*) expression, TE methylation level, global mCG level, and epimutation rates across RILs. *VIM2/4* expression is significantly negatively correlated with mCG level at AT1TE80755 (**b**) and with global mCG levels (**c**), and positively correlated with global CG epimutation rates (calculated as 0.5×(α+β)) (**d**). Linear regression lines (blue), 95% confidence intervals (shaded area), and statistics (Pearson’s R, p-values) are shown. Together, these results indicate that in the Cvi genotype, elevated *VIM2/4* expression is associated with reduced methylation levels and higher epimutation rates. Two extreme *VIM2/4* expression data points were outliers and removed for linear regression fitting. **e**, PCR validation of the Cvi-specific deletion in selected RILs confirms genotype-specific segregation. **f**, qPCR-based quantification of combined *VIM2* and *VIM4* (i.e., *VIM2/4*) expression in selected RILs shows obvious elevated expression in lines carrying the Cvi-specific deletion. Expression is normalized to Col-0 as a reference. **g**, Gene co-expression correlation network for key methylation pathway genes, comparing RILs with Ler-derived (PeakLer, left) and Cvi-derived (PeakCvi, right) alleles at the chromosome 1 QTL. The dashed box highlights strong co-expression between *VIM2/4* and core components of the CG methylation maintenance machinery, including *MET1*, *VIM1*, and *VIM3*. Notably, this module also includes *CMT3*, a key CHG methyltransferase, suggesting coordination between CG and non-CG methylation pathways. The solid box indicates altered co-expression between methylation maintenance genes and DNA demethylases (*DML2-3*, *ROS1*, *DME*), reflecting network-level dysregulation between methylation and demethylation pathways in the Cvi genotype. The presence of the Cvi-specific deletion disrupts co-expression patterns within the methylation regulatory network, particularly diminishing *VIM2/4* connectivity. Heatmap values represent Pearson’s correlation coefficients. **h**, Expression levels of DNA methylation regulators in PeakLer and PeakCvi lines. Genes with significant expression differences between genotypes are highlighted. Error bars represent SEM; p-values from two-sided t-tests (*p < 0.05, **p < 0.01).

To confirm the structural variant in the MAML population, we performed PCR genotyping in selected RILs, which revealed genotype-specific segregation of the deletion (Fig. 4e and Extended Data Fig. 3a). We then quantified the combined expression levels of *VIM2* and *VIM4* (i.e., *VIM2/4*) by qPCR and observed a consistent increase in expression in deletion-containing lines (Fig. 4f and Extended Data Fig. 3b). Notably, the expression level of *VIM2/4* was highly predictive of both mCG levels and spontaneous epimutation rates in global and gbM regions (Fig. 4c,d and Extended Data Fig. 3c,d), pointing at a strong functional impact on epigenomic instability. On average, the Cvi-specific deletion reduces global mCG levels by 1.7% and gbM levels by 5.5%, and is associated with a ∼1.5-fold increase in epimutation rates genome-wide and ∼1.9-fold within gbM regions (Fig. 4c,d and Extended Data Fig. 3c,d).

Although elevated *VIM* expression might intuitively be expected to promote methylation maintenance, our findings suggest the opposite. A previous study has shown that experimental overexpression of *VIM1* or *VIM3* leads to hypomethylation ^12,38^, indicating that maintenance of CG methylation requires tightly regulated *VIM* dosage. The mechanistic basis remains unclear, but overexpression has been proposed to disrupt stoichiometry within methylation maintenance complexes or to exert dominant-negative effects, possibly through sequestration of essential cofactors ^38,39^. Given the high sequence similarity among *VIM* gene family ^12,38^, a similar mechanism may apply to *VIM2* and *VIM4*.

To investigate how variation in *VIM* dosage influences broader epigenetic regulatory networks, we examined gene co-expression patterns across the population. In both the MAML population and natural accessions, *VIM2* and *VIM4* showed strong co-expression not only with each other but also with core components of the CG methylation machinery, including *MET1*, *VIM1*, *VIM3*, and the CHG maintenance methyltransferase *CMT3* (Fig. 4g and Extended Data Fig. 3e), suggesting their integration into the central DNA methylation network. Notably, this co-expression architecture was disrupted in RILs carrying Cvi-derived alleles, particularly for *VIM2/4*, whose connectivity with methylation-related genes—including DNA demethylases—was markedly reduced (Fig. 4g,h). In particular, the demethylases *ROS1* was significantly downregulated in RILs carrying the deletion (Fig. 4h). This network-level dysregulation likely contributes to the elevated rates of both stochastic mCG losses and gains seen in the Cvi QTL background.

### QTL predicts increased epigenetic drift and functional divergence

Our findings in the MAML population predict that natural accessions that are fixed for the Cvi-derived deletion should exhibit not only reduced global CG methylation, but also increased methylation variance due to epigenetic drift. Over evolutionary timescales, such drift could drive progressive methylation divergence between lineages that carry the Cvi-derived haplotypes—a possibility not examined in previous studies of natural accessions. To test this, we re-analyzed DNA methylome data from the natural Cvi accessions population described by Zicola et al. (co-submission). Accessions carrying the deletion between *VIM2* and *VIM4* showed significantly lower global mCG levels and a clear trend toward increased methylation variance compared to those without the deletion (Fig. 5a,b). To ask whether similar patterns are detectable more broadly, we analyzed publicly available SNPs located within the Cvi-specific deletion interval across natural accessions. Although the full structural variant is not directly genotyped in these data, we used these SNPs as haplotype proxies. Consistent with our predictions, the minor allele was significantly associated with both reduced global mCG levels and increased methylation variance across natural accessions (Fig. 5c,d). We reasoned that the QTL-induced increase in epigenetic drift could also drive functional divergence at the chromatin level. To assess this experimentally, we performed ATAC-seq in two RILs with contrasting spontaneous CG epimutation rates and tracked chromatin accessibility dynamics across multiple generations (Supplementary Fig. 2). Notably, the RILs carrying the Cvi genotype at the chromosome 1 QTL, and exhibiting higher epimutation rates, also showed visibly greater divergence in accumulation of chromatin accessibility patterns than the low-rate line (Fig. 5e; Methods). This indicates that reduced CG maintenance fidelity correlates with functional divergence over generations.

**Fig. 5.**
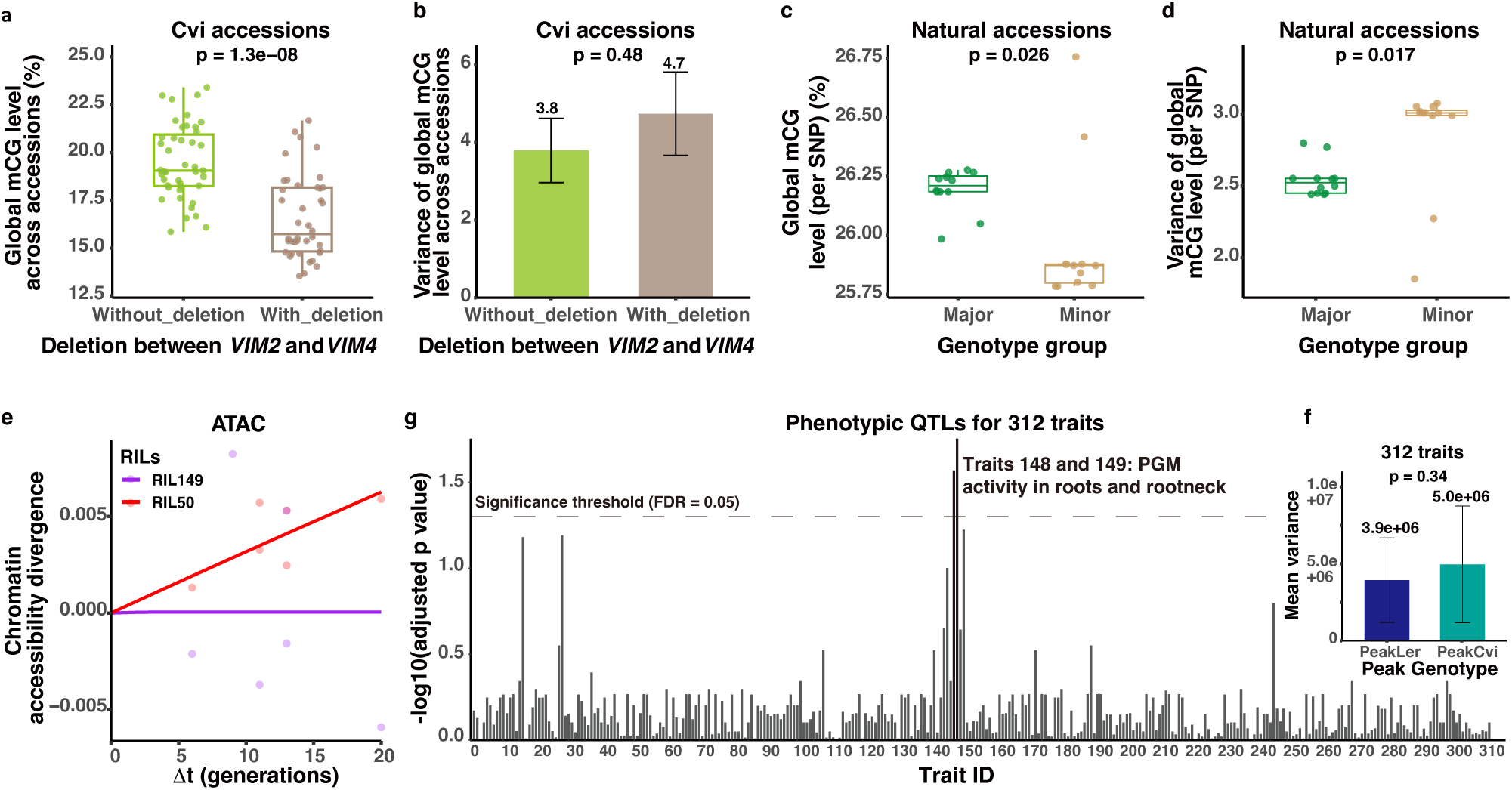
Epigenomic, chromatin, and phenotypic consequences of the Chr1 QTL and Cvi-specific deletion. **a–b**, Global mCG levels (**a**) and variance (**b**) in Cvi accessions stratified by presence or absence of the deletion between *VIM2* and *VIM4*. Deletion-carrying accessions exhibit significantly lower methylation levels (two-sided t-test) and a trend toward increased variance (F-test). **c–d**, Global mCG levels (**c**) and variance (**d**) across natural accessions grouped by SNP genotypes within the Cvi-specific deletion interval. Accessions carrying the minor allele—used as a proxy for the deletion—exhibit significantly reduced mCG levels and increased methylation variance (two-sided t-test). **e**, Divergence in chromatin accessibility over generations in two RILs with contrasting epimutation rates. The hyper-epimutable line (RIL50, Cvi genotype at the Chr1 QTL) exhibits visibly greater accumulation of chromatin accessibility divergence over time compared to the stable line (RIL149), indicating that reduced epigenetic fidelity is coupled to functional divergence. Estimated rates of chromatin accessibility changes are substantially elevated in RIL50 (gain rate: 1.7 × 10 ⁻⁴; loss rate: 2.8 × 10⁻³) relative to RIL149 (gain rate: 1.3 × 10⁻⁹; loss rate: 2.4 × 10⁻⁸). **f**, Phenotypic variance across 312 traits in the Ler × Cvi RILs population. RILs carrying the Cvi allele at the chromosome 1 QTL peak show increased trait variance on average. P-values were calculated using a paired two-sided t-test. **g**, Phenotypic QTL (pQTL) mapping results for 312 traits. Traits 148 and 149 (phosphoglucomutase (PGM) activity in roots and root neck) show significant associations with the Chr1 meQTL, suggesting downstream phenotypic consequences of altered methylation dynamics. The dashed line indicates the False Discovery Rate (FDR) threshold (adjusted p = 0.05, calculated using the Benjamini-Hochberg procedure).

The dual impact of the chromosome 1 QTL on mCG levels, epimutation rates, and functional variation raises the possibility that it influences downstream phenotypes, either by increasing trait variance through epigenetic drift or by affecting trait means through shifts in mCG levels. We found evidence supporting both mechanisms. Specifically, we leveraged previously reported phenotypic QTL (pQTL) data encompassing 312 traits measured in the Ler × Cvi RILs population ^29,30^. On average, the Cvi genotype exhibited higher phenotypic variance across all traits (Fig. 5f). In addition, two traits corresponding to phosphoglucomutase (PGM) activity in roots and root neck (traits 148 and 149) showed significant associations with the major-effect methylation QTL on chromosome 1 (Fig. 5g, Extended Data Fig. 4a, and Supplementary Table S11). All three *PGM* genes exhibited hypomethylation in lines with the Cvi genotype (Extended Data Fig. 4b,c). This links *VIM2/4* overexpression to altered PGM enzymatic activity and root growth inhibition, consistent with prior findings that *VIM1/3* overexpression reduces DNA *methylation* and inhibits root growth ^12,38,39^. Careful phenotypic surveys combined with re-sequencing of the *VIM2/4* regions should be carried out in natural populations to determine whether the observed patterns reflect increased phenotypic plasticity and potential adaptive advantages in dynamic environments.

## Discussion

Our study identified a major-effect QTL in *Arabidopsis thaliana* that jointly controls CG methylation levels and spontaneous epimutation rates (Figs. 2 and 3). This QTL is caused by a naturally occurring deletion between *VIM2* and *VIM4*, which disrupts the expression of these core DNA methylation regulators as well as other co-expressed methylases and demethylases ^1,12,13,38–40^ (Fig. 4). We showed that this disruption causes a genome-wide shift to lower mCG levels, accompanied by increased methylation gain and loss rates—leading to accelerated epigenetic drift and functional divergence within genotypes over generations (Figs. 2-5). These findings help explain why the Cvi population, which recently colonized the Cape Verde Islands and experienced a strong bottleneck approximately 5,000 years ago ^41^, has undergone rapid epigenomic divergence—potentially driven by the *VIM2/4* deletion. Interestingly, the QTL had its strongest effect on gbM genes (Figs. 2 and 3)—regions typically regarded as epigenetically stable and, more recently, also used in epigenetic clock models to estimate evolutionary divergence ^21^. Our findings here suggest that such clocks are only reliable when the DNA methylation maintenance machinery is sufficiently intact.

An earlier attempt to identify genetic modifiers of gbM used F2 intercrosses and assessed somatic deviations from expected methylation patterns as indirect indicators of epimutation ^18^. Although informative, this approach is limited in its ability to draw definitive conclusions about the dynamics and mechanisms of epimutations across generations. In contrast, our multigenerational experimental design has revealed a two-step “punctuated equilibrium” model of mCG evolution. In this model, genetic perturbation causes a rapid shift in methylation steady-state, followed by a secondary phase in which reduced maintenance fidelity leads to elevated epimutation rates and gradual epigenetic divergence ^36^ (Figs. 1-5). This coupling of fast and slow processes offers a framework for how genetic and epigenetic factors interact to shape heritable variation. These insights also have practical implications. Some genotypes may be predisposed to higher rates of epigenetic drift, influencing the emergence of epigenetic variation in natural populations and potentially complicating efforts to maintain stable germplasm in breeding programs. Future work should explore whether similar variation in methylation fidelity occurs across other genotypes, species, or environmental conditions.

Understanding how genetic modifiers influence epigenetic stability could open new avenues for crop improvement by balancing adaptive epigenetic flexibility with the need to preserve functional genome integrity. It may be interesting to perform selective breeding for high or low epimutation rates in various plant species to examine if this leads to specific phenotypic trade-offs. Such an approach may reveal general insights into the evolutionary optimization of the CG methylation maintenance system itself.

## Methods

### WGBS and transcriptome data of natural, MA, and Cvi accessions

This study integrates newly generated and previously published data from four major sources: natural accessions, MA-Accessions, Cvi accessions, and Mutation Accumulation Mapping Lines (MAML) Population.

Raw WGBS data for 927 samples were downloaded from the NCBI Sequence Read Archive (SRA; accession SRP018263), corresponding to GEO series GSE43857 ^4^. The corresponding transcriptome data were obtained from GSE80744 ^4^. SRA files were converted to FASTQ format using prefetch (v2.9.2). For samples with multiple FASTQ files associated with the same GSM accession, individual BAM files were first generated and then merged using “samtools merge -n” (Samtools v1.11). Only samples derived from leaf tissue were retained. To account for redundant accessions, we selected the sample (GSM ID) with the highest sequencing depth when multiple samples corresponded to the same ecotype ID. This filtering resulted in a final set of 850 unique accessions (Supplementary Table 1).

To begin to quantify the impact of genetic variation on epimutational processes in plants, we previously generated mutation accumulation (MA) lines using 4 different *A. thaliana* natural ecotypes as founders (i.e., MA-Accessions, including Col-0, Kn-0, Mt-0, and Tsu-0) ^21^. These lines were propagated under uniform conditions for 16 generations (G16) (Fig. 1b). The founder ecotypes were carefully selected to represent the broad range of genome-wide CG methylation levels that have been observed in a species-wide methylome survey of *A. thaliana* (Fig. 1a) ^2,4,14^. Further details concerning the construction of these pedigrees and DNA extraction protocols are provided in the original publication ^21^.

Additional WGBS data from Cvi accessions were obtained from Zicola et al. (co-submission).

### Construction of Mutation Accumulation Mapping Lines (MAML) population

To be able to identify specific genetic loci contributing to intra-specific variation in epimutation rates, in this study, we newly generated a unique selfing-derived MAML in the model plant *A. thaliana* (Fig. 1f). The MAML comprise 68 genetically distinct MA pedigrees (with eight lineages each), which are derived from 68 different founders Recombinant Inbred Lines (RILs). These RILs were initially obtained from a cross between Landsberg erecta (Ler-0) and Cape Verde Islands (Cvi-0) accessions. A key advantage of using the Ler x Cvi RILs panel as MA-pedigree founders is that these lines have well-defined genetic backgrounds, and that the G0 founders (or siblings thereof) have been subject to previous QTL mapping studies involving hundreds of classical phenotypes as well as high-throughput molecular traits, such as mRNA, metabolite and protein abundance ^29,30^. In addition, the Ler-0 and Cvi-0 genotypes have significantly different epimutation rates (Fig. 3a), and Cvi has unusually low genome-wide methylation levels (Fig. 1a), making it an ideal crossing partner to assess haplotype-specific epimutation dynamics in otherwise recombinant Ler and Cvi genomes. The MAML constitute the first experimental system to study the genetic basis underlying epimutational processes in plants.

The MAML were grown in climate-controlled chambers for ten selfing generations (long-day conditions, 16-h day lengths, 18 °C). Seeds from each generation have been stored. To obtain plant material for WGBS, we grew plants from G2 and G9 seeds, following the sampling scheme shown in Fig. 1f. Within each pedigree, this sampling scheme gives us divergence times (Δt) between samples ranging from 2 to 18 generations (Fig. 3b), which is sufficient for epimutation rate estimation. Samples were collected from young above-ground tissue. Leaf tissue was flash frozen in liquid nitrogen, and DNA was extracted using a Qiagen Plant DNeasy kit (Qiagen, Valencia, CA, USA) based on the manufacturer’s instructions. MethylC-seq libraries were prepared based on the protocol described in Urich et al. (2015) ^42^. Libraries were sequenced at Genewiz on a NovaSeq 6000 platform (Illumina) in a paired-end 150 bp format.

### WGBS data processing

To ensure consistent processing across multiple *Arabidopsis* WGBS datasets, including our MAML data, natural accessions, and MA-accessions, we employed the MethylStar pipeline ^43,44^ with TAIR10 ^45^ as the reference genome. MethylStar is a fast and robust pre-processing pipeline for bulk WGBS data ^44^. Cytosine-level methylation states were inferred using Methimpute ^43^, which classifies each site into one of three states: Unmethylated (“U”), Intermediate (“I”), or Methylated (“M”), based on a three-state Hidden Markov Model. To ensure high-quality methylation state calls, only cytosines with a maximum posterior probability larger than 0.99 were retained for downstream epimutation analysis. Sequencing metrics for each MAML sample, including sequencing depth, mapping efficiency, and coverage, are provided in Supplementary Table 2.

### Enrichment of annotations

Gene and transposable element (TE) annotation files in GFF3 format were downloaded from Ensembl Plants (http://plants.ensembl.org/info/data/ftp). The gbM, teM, and LM gene sets were obtained from previous work ^46,47^. In this study, genes previously labeled as unmethylated (UM) were referred to as lowly methylated (LM), following Zhang et al. (2024) ^48^. The “Other_gene” category was defined by subtracting gbM, teM, and LM genes from the total gene set. Intergenic regions were defined as genomic intervals not overlapping with genes, TEs, or promoter regions, using bedtools complement ^36,49^. The “Others” category includes genomic regions not assigned to any of the above annotations. The genome coordinates and corresponding epigenetic marks for the 36 chromatin states (CSs) were obtained from the PCSD database ^37^. Clustering of these CSs into four major groups was performed according to Hazarika et al. ^36^, corresponding to green, red, blue, and purple states.

### Calculation of average methylation levels

Average observed methylation levels were computed using methylome data, considering only sites with posteriorMax ≥ 0.99 to ensure high-confidence calls. The proportions of unmethylated (U) and intermediate (I) states were calculated relative to the total number of high-confidence sites (n_total). The methylation level was then defined as “1 - nU/n_total - 0.5 × nI/n_total”, where nU and nI represent the number of sites classified as unmethylated and intermediate states, respectively. Predicted methylation levels were obtained by applying the mathematical model formulated by van der Graaf et al. (2015) ^9^, which relates steady-state methylation to the gain and loss rates.

### Estimation of epimutation rates

To obtain accurate estimates of epimutation rates in the MAML pedigrees, global and specific annotated estimates of the CG epimutation gain rate (α) and the loss rate (β) at the level of individual cytosines were acquired using the R package AlphaBeta (version 1.10.0) ^22^. AlphaBeta is a generalization of our previous modeling-based approach for estimating epimutation rates in selfing-derived MA lines ^9^. AlphaBeta takes pedigree-based multi-generational WGBS data as input and outputs estimates of the unknown gain (α) and loss rates (β), along with their confidence intervals. To quantify CG methylation divergence between samples, we applied the divergence estimation function implemented in the AlphaBeta package, which leverages pedigree structure to model changes in methylation states. The underlying methodology and mathematical framework are detailed in the original publication ^22^. For all analyses, we fitted the neutral model (“ABneutral”).

For comparison, we also applied a count-based method to estimate epimutation rates on MAML. Our pipeline categorizes methylation states into three types: Methylated (M), Unmethylated (U), and Intermediate (I). To quantify epimutation rates, we tracked methylation state transitions between early and late generations (i.e., G2 and G9) within the main lineage. Methylation gain was assessed by calculating the ratio of U to M transitions, defined as the number of sites that changed from U to M relative to the total number of U sites in the early generation, and the ratio of U to I transitions, representing the proportion of U sites that became I. Similarly, methylation loss was quantified by the ratio of M to U transitions, calculated as the number of sites transitioning from M to U relative to the total M sites in the early generation, and the ratio of M to I transitions, reflecting the proportion of M sites that shifted to I. We highlight the advantages of the AlphaBeta computational framework over the count-based approach in capturing epimutation rates in Supplementary Fig. 3.

### Identification of SNP markers from WGBS for genetic mapping

To construct a genetic map for Ler x Cvi RILs using single-nucleotide polymorphism (SNP) markers, we identified polymorphic SNPs from WGBS data of MAML. BAM files were first extracted from the bismark-deduplicate output generated by Methylstar ^43,44^, followed by sorting and indexing with samtools (version 1.17). Genome-wide SNPs were identified using bcftools (version 1.17) with default settings, using TAIR10 as the reference genome while excluding indels.

To ensure the selected SNPs were informative for genetic mapping in Ler x Cvi RILs, we cross-referenced SNPs identified from WGBS with polymorphisms between Ler-0 and Cvi-0 obtained from the 1001 *Arabidopsis* Genomes project ^50^. Only SNPs that overlapped between these datasets were retained. As a result, the set of SNPs consisted exclusively of two genotypes corresponding to Ler and Cvi.

Further filtering was applied to select homozygous SNPs and retain only SNPs with a single nucleotide in both alleles to exclude complex variants. Next, consensus SNPs were identified across all samples within the same RIL to ensure consistency, followed by additional filtering to retain SNPs with an occurrence frequency between 30% and 70% across all RILs, thereby excluding rare or nearly fixed variants. To minimize sequencing artifacts inherent to WGBS, we restricted the selected SNPs to A/G transitions. Since WGBS induces C-to-T conversions due to bisulfite treatment, C/T SNPs can be confounded with methylation status rather than true genetic polymorphisms. By focusing on A/G SNPs, we aimed to reduce the likelihood of misidentifying methylation-associated changes as genetic variants. After applying these filtering criteria, we obtained a set of 63,359 SNPs.

To reduce redundancy and noise in SNP markers, we applied a sliding-window approach with a 200-SNPs window size and a 5-SNPs step size. Within each window, the SNP classification was determined based on the proportion of variant sites. Windows where more than 50% of the SNPs were classified as Ler were assigned as Ler markers, whereas those with fewer than 50% Ler SNPs were assigned as Cvi markers. If the proportion of Ler and Cvi SNPs was exactly 50%, the window was assigned as missing data (NA). Applying this classification, we obtained 12,479 SNP markers from the sliding-window analysis.

Further refinement was performed using the qtl R package ^51^, where redundant markers were removed with the “findDupMarkers” function, and markers with Ldiff > 2 were excluded using the “droponemarker” function to improve map quality. This resulted in a final set of 732 SNP markers (Supplementary Table 4), which were subsequently used to construct a high-density genetic map. Comparison with 144 physical markers ^29^ confirmed the reliability of these SNP markers, significantly improving the accuracy of the genetic map (Fig. 1g, Supplementary Fig. 1, and Supplementary Table 3), which in turn enabled higher-resolution QTL mapping and more precise localization of candidate QTL regions.

### QTL mapping approach

Using methylation levels as molecular quantitative traits, we performed methylation QTL (meQTL) for CG, CHG, and CHH context-specific methylation levels separately for genome-wide as well as annotation-specific methylation levels (Fig. 2). We also tested meQTL for methylation level at early generation (i.e., G2) and late generation (i.e., G9) separately (Extended Data Fig. 2a). Treating the genotype-specific estimates for the gain (α) and loss rate (β) as molecular quantitative traits, we performed a search for QTL (i.e., αβ-QTL) underlying rate variation using classical linkage analysis. QTL scans were performed for genome-wide as well as annotation-specific epimutation rates (Fig. 3).

Previous studies have used the Ler x Cvi RILs panel to perform system-wide QTL analysis of population-level transcriptomes, metabolomes, and proteomes ^29,30^. They applied QTL scans to 24,065 transcript abundance traits (local and distant eQTL), for 2,843 protein abundance traits, and for 312 phenotypic traits (pQTL). We re-analyzed these molecular data and searched for specific linkage association with our αβ-QTL and meQTL. This approach can uncover genes and their products that act in candidate pathways or co-expression networks ^52^, and should provide experimental targets for follow-up molecular work. Due to the high sequence similarity and close physical proximity (∼2.6 kb apart) between the paralogous genes *VIM2* and *VIM4*, their transcripts are difficult to reliably distinguish by RNA-seq. As a result, the raw eQTL dataset reports expression values only for *VIM4*, which likely reflect the combined expression of both *VIM2* and *VIM4* (hereafter referred to as *VIM2/4*).

QTL mapping was performed using the qtl R package to identify loci associated with epigenetic variation ^51^. The genetic map was constructed using est.map(), incorporating an optimized error probability of 1 × 10⁻⁷ for improved accuracy. Single-QTL genome scans were conducted using Haley-Knott regression for multiple phenotypic traits, including global and annotation-specific methylation levels and epimutation rates. Significance thresholds (α = 0.05) were determined via 1000 permutations. LOD Confidence Intervals (CIs) were defined using a drop threshold of 2.0 via the “lodint()” function, and the most significant QTL peak LOD for each trait was identified.

### Call Cvi- and Ler-derived chromosomal segments

To define genomic segments inherited by the Ler x Cvi RILs, we classified each genomic region into Ler segments and Cvi segments based on the previously mentioned 12,479 SNP markers, which exclusively comprised Cvi and Ler genotypes. For each RIL, markers assigned to Ler were grouped into contiguous segments (Supplementary Fig. 1). The start and end positions of each Ler segment were determined using the first and last marker of the segment, with the midpoint taken as the representative position. Cvi segments were identified independently using the same approach, based on markers assigned to Cvi.

### ATAC-seq and data processing

ATAC-seq was performed across generations in two RILs with contrasting epimutation rates— RIL50, a hyper-epimutable line carrying the Cvi genotype at the Chr1 QTL, and RIL149, a stable line with the Ler genotype at the same locus. ATAC-seq samples were collected from the immediate progeny of the sibling plants used for WGBS, ensuring temporal and genetic continuity between chromatin accessibility and methylation measurements. (Supplementary Fig. 2).

The *Arabidopsis thaliana* RILs were grown under long-day conditions (16 h light / 8 h dark) on ½ MS plates at 25 °C for 7-9 days. For the nuclei extraction, approximately 200 mg of shoots were collected. The samples were chopped down in 1 ml pre-chilled lysis buffer (10 mM MES-KOH pH5.4, 10 mM NaOH, 250 mM sucrose, 0.1 mM spermine, 0.5 mM spermidine, 1 mM DTT, 1% BSA, 0.5% TritonX-100) and filtered through a 40 µm and a 20 µm strainer, with a centrifugation step in between (500 xg, 5 min). The nuclei were finally pelleted with a gradient solution (35% Percoll in lysis buffer) in a swing centrifuge at 500 xg for 10 min and resuspended in tagmentation buffer (10 mM TAPS-NaOH pH 8.0, 5 mM MgCl2). Nuclei quality and concentration were checked with DAPI staining at a cell counter. 50k - 100k purified nuclei were incubated with 3 µl Tn5 enzyme in 40 µl tagmentation buffer (s.o.) at 37 °C for 30 min. The products were purified using a PCR-extraction kit (NEB Monarch PCR & DNA Clean-up, #T1030) and then amplified for 10-12 cycles using Q5 polymerase (NEBNext HiFi 2x PCR Mastermix, #M0541). PCR conditions were followed as described previously ^53^. Amplified libraries were purified with AMPure beads, and concentrations were determined using Qubit and qPCR (Roche KAPA Library Quantification Kit) prior to sequencing. Paired-end 150 bp sequencing was performed on an Illumina NovaSeq 6000 platform.

Sequencing reads were processed using a standard ATAC-seq pipeline as described below. Adapters were trimmed using Trimmomatic v0.39, and paired-end reads were aligned to the TAIR10 genome using BWA-MEM (v0.7.17). Sorted BAM files were deduplicated using Picard MarkDuplicates (v2.23.8; REMOVE_DUPLICATES=true). Peaks were called with MACS2 (v2.2.9.1) using BEDPE input mode and *Arabidopsis* genome size (1.35e8), with parameters -- keep-dup all, --SPMR, and -q 0.05. All downstream analyses were restricted to chromosomes 1– 5.

To quantify chromatin accessibility divergence across generations, we converted ATAC-seq peak calls into a genome-wide binary presence/absence matrix. The *A. thaliana* genome was divided into non-overlapping 100 bp bins. Each bin was scored as “M” (accessible) in a sample if at least 50% of its length overlapped with a called ATAC-seq peak, and as “U” (inaccessible) otherwise. This binarization approach mirrors the construction of the methylation matrix from WGBS data. All bins were retained regardless of accessibility to ensure full genome coverage. Chromatin accessibility divergence between samples across generations, as well as gain and loss rates of accessibility states, were inferred using AlphaBeta ^22^, a pedigree-based model originally developed for estimating epimutation dynamics from WGBS data.

### PCR genotyping and qPCR

For extraction, each DNA (Qiagen DNeasy Plant Pro Kit, #69206) and RNA (NEB Monarch Total RNA Miniprep Kit, T2010) sample was taken from leaves of the same plant, after growing for 4 weeks on soil (16 h light / 8 h dark; 23 °C; 70% RH). Samples were frozen in liquid N₂ and stored at –80 °C for 1–3 days prior to extraction, except for RNA from RIL162, which was stored for 3 weeks due to re-extraction from a new leaf following initial failure. Homogenization was done mechanically in a bead mill (QIAGEN, TissueLyser II).

The genotype was determined by PCR (Tm=65°C, 30 cycles) using OneTaq DNA Polymerase (NEB, #M0480) and visualised on a 1% agarose gel. RNA extraction yield was measured with spectrophotometry (DeNovix, DS-11 FX+) and resulted in 4.5 µg – 13.2 µg of total RNA with A260/A280 ratios of 2.06 – 2.28. For cDNA synthesis (Thermo Scientific RevertAid First Strand cDNA Synthesis Kit, #K1621), 1 µg of DNase-treated total RNA was used. mRNA was selected by using Oligo(-dT)-primer.

qPCR was performed using Luna Universal qPCR MasterMix (NEB, #M3003S) on a Roche LightCycler 480 (Software version 1.5.0) with three technical replicates for each sample. *ACTIN2* (AT3G18780) was selected as an internal control after testing three potential control genes based on the least dispersion between samples (SD=0.44). The primer pair for ACT2 produces a single amplicon of 179 bp with a tested efficiency of 111%. The primer pair for *VIM2*/*4* produces a single amplicon of 115 bp with a tested efficiency of 110%. Following the Luna Protocol, the qPCR run used a 20 µl reaction with 1 µl of 1:5 diluted cDNA as template and was performed for 45 cycles. NT-control for all runs resulted in CT values of 34.62 (SD=1.2) for *VIM* expression and 36.44 (SD=0.51) for ACT2 expression. The transcript levels of the target genes were quantified with comparative CT-method normalized to AtACTIN2 in Col-0. Due to the high sequence similarity and close physical proximity (∼2.6 kb apart) between the paralogous genes *VIM2* and *VIM4*, their transcripts are difficult to reliably distinguish by qPCR. Therefore, our qPCR assay captures the combined expression level of both *VIM2* and *VIM4* (i.e., *VIM2/4*).

All Oligo sequences are listed below:

Genotyping: gDEL2778-F: GCATAAATTAGCGAGTGAAAACGAACCGTGACAG gDEL2778-R: GCATAAATTAGCGAGTGAAAACTAACCGTAACAGG
qPCR: qVIM-F: GCTGAAGTTGCGGAGTCATCAAAC qVIM-R: CTGAGAGACTTGTACCACCAACAGAG qACT2-F: GTTCCAGCCCTCGTTTGTG qACT2-R: CAAGTGCTGTGATTTCTTTGCTC

## Acknowledgements

We thank Ioanna Kakoulidou, Maria-Cecília D. Costa, members of R.J.S.’s laboratory, and the TUM Duernast community for their valuable technical assistance. We also thank Joost J. B. Keurentjes for generously sharing seeds and the eQTL and pQTL data for the Ler × Cvi RILs. F.J. acknowledges funding from the Deutsche Forschungsgemeinschaft (DFG) under project number 460890770. Z.Z. holds a fellowship from the China Scholarship Council (no. CSC202006380020). This research was supported by the National Science Foundation (MCB-2242696) to R.J.S.

## Author contributions

F.J. conceived and designed the study. Z.Z. analyzed data. F.J., R.J.S., W.W., and Y.X. generated WGBS and ATAC-seq data. W.W. performed PCR and qPCR experiments. J.Z. and A.M.H. generated and provided Cvi accessions data. Z.Z. and F.J. wrote the manuscript, with contributions from all authors.

## Ethics declarations

### Competing Interests

The authors declare no competing interests.

## Notes

### Competing Interest Statement

The authors have declared no competing interest.

